# Seasonal dynamics of a glycan-degrading flavobacterial genus in a tidally-mixed coastal temperate habitat

**DOI:** 10.1101/2023.03.30.534869

**Authors:** Maéva Brunet, Nolwen Le Duff, Fabienne Rigaut-Jalabert, Sarah Romac, Tristan Barbeyron, François Thomas

**Author notes:** Address correspondence to François Thomas.

## Abstract

Coastal marine habitats constitute hotspots of primary productivity. In temperate regions, this is due both to massive phytoplankton blooms and dense colonization by macroalgae that mostly store carbon as glycans, contributing substantially to local and global carbon sequestration. Because they control carbon and energy fluxes, algae-degrading microorganisms are crucial for coastal ecosystem functions. Environmental surveys revealed consistent seasonal dynamics of alga-associated bacterial assemblages, yet resolving what factors regulate the *in situ* abundance, growth rate and ecological functions of individual taxa remains a challenge. Here, we specifically investigated the seasonal dynamics of abundance and activity for a well-known alga-degrading marine flavobacterial genus in a tidally-mixed coastal habitat of the Western English Channel. We show that members of the genus *Zobellia* are a stable, low-abundance component of healthy macroalgal microbiota and can also colonize particles in the water column. This genus undergoes recurring seasonal variations with higher abundances in winter, significantly associated to biotic and abiotic variables. *Zobellia* can become a dominant part of bacterial communities on decaying macroalgae, showing a strong activity and high estimated *in situ* growth rates. These results provide insights into the seasonal dynamics and environmental constraints driving natural populations of alga-degrading bacteria that influence coastal carbon cycling.

**Originality-significance statement:** Glycan-degrading bacteria play a crucial role in marine habitats to remineralize organic carbon sequestered in algal biomass. Yet, resolving what factors regulate the *in situ* abundance, growth rate and ecological functions of individual taxa remains a challenge. Here, we investigate the seasonal dynamics of abundance and activity of an environmentally relevant glycan-degrading bacterial genus in two constrasted compartments of the same coastal habitat, i.e. the surface of diverse macroalgae and the water column. These results provide insights into the recurring temporal patterns and environmental constraints driving natural populations of alga-degrading bacteria that influence ocean carbon cycling.

## Introduction

Coastal marine habitats are unique and dynamic ecosystems at the interface between continents and seas. They constitute hotspots of primary productivity, accounting for ca. 20% of the global primary production on Earth (Gattuso *et al*., 1998). In temperate regions, this is largely due both to massive phytoplankton blooms and dense colonization by macroalgae that mostly store carbon in the form of glycans, contributing substantially to local and global carbon sequestration (Santos *et al*., 2022). Microbial breakdown represents a crucial bottleneck to re-inject the large pool of algal organic matter in the marine carbon cycle (Buchan *et al*., 2014), by (i) making it accessible for higher trophic levels, (ii) liberating dissolved or particulate matter for local recycling or further export, (iii) remineralizing it back to atmospheric CO2 and (iv) eventually influencing how much carbon is sequestered. Because they control the fluxes of carbon and energy both locally and between adjacent zones, algae-degrading micro-organisms are of critical importance for coastal ecosystem functions. The role of microorganisms as recyclers of algal biomass is even more relevant in the context of global change, since their action can influence the carbon balance in coastal ecosystems already threatened by human activities and pollution.

Surveys of natural coastal microbial communities have revealed the major role of heterotrophic bacteria in processing algae-derived compounds. Metagenomic analyses of long-term bacterioplankton series notably show predictable substrate-controlled successions of distinct bacterial taxa following phytoplankton blooms, mainly belonging to *Flavobacteriia*, *Gammaproteobacteria* classes and to the genus *Roseobacter* within *Alphaproteobacteria* (Gilbert *et al*., 2009; Teeling *et al*., 2012, 2016). Macroalgae also host abundant and diverse epibiotic bacterial communities (Egan *et al*., 2013; Martin *et al*., 2014; Singh and Reddy, 2016), the composition of which can greatly vary depending on algal species, thallus part, seasons and locations. Similar to phytoplankton bloom demise, breakdown of soluble macroalgal glycans and intact tissues is also accompanied by a succession of distinct heterotrophic bacterial clades (Enke *et al*., 2019; Bunse *et al*., 2021; Brunet *et al*., 2021). Environmental surveys revealed consistent seasonal dynamics of alga-associated bacterial assemblages, yet resolving what factors regulate the *in situ* abundance, growth rate and ecological functions of individual taxa remains a challenge (Bunse and Pinhassi, 2017). Studies on cultivated strains provided detailed insights into the metabolic pathways marine heterotrophic bacteria use to degrade algal biomass (Thomas *et al*., 2012; Kabisch *et al*., 2014; Ficko-Blean *et al*., 2017; Koch *et al*., 2019; Reisky *et al*., 2019; Sichert *et al*., 2020), but do not inform on the influence of biotic interactions with their host and abiotic constraints, hindering our understanding of their ecological impact.

Here, we specifically investigated the seasonal dynamics of abundance and activity for an alga-degrading marine flavobacterial genus in a tidally-mixed coastal habitat of the Western English Channel, an epicontinental sea with strong seasonal and interannual variations of climatic, hydrological and biological conditions (Goberville *et al*., 2010; Lefran *et al*., 2021). We analyzed two complementary time-series, including (i) a yearlong monthly epibiota sampling on 5 different healthy macroalgal species and decaying tissues and (ii) bi-weekly water samples over an 8-year period. Using (RT-)qPCR, we targeted the genus *Zobellia*, which has received much attention in the last two decades as a specialist of macroalgal degradation among *Flavobacteriia* (Lami *et al*., 2021). Six out of the eight validly described *Zobellia* species were previously isolated from algal surfaces and the others from beach sand and seawater (Barbeyron *et al*., 2001, 2021; Nedashkovskaya *et al*., 2004; Nedashkovskaya *et al*., 2021). In addition, *Zobellia* genes were reported in molecular studies of macroalgal microbiomes (Miranda *et al*., 2013; Dogs *et al*., 2017) and coastal seawater (Alonso *et al*., 2007). *Zobellia* members are known to be efficient degraders of algal purified polysaccharides (Thomas *et al*., 2017) as well as fresh algal tissues (Brunet *et al*., 2022).

Hence, the genus *Zobellia* was reported to play a key role in algal biomass recycling but there is currently no data regarding its natural distribution. In the present study, we show that the genus *Zobellia* is a stable, low-abundance component of healthy macroalgal microbiota and can also be found mostly associated to particles in the water column. It undergoes recurring seasonal variations with higher abundances in winter, significantly associated to biotic and abiotic variables. *Zobellia* spp. can become a dominant part of bacterial communities on decaying macroalgae, showing a strong activity and high estimated *in situ* growth rates. These results provide insights into the seasonal dynamics and environmental constraints driving natural populations of alga-degrading bacteria that influence coastal carbon cycling.

## Experimental procedure

### Sampling

Surface microbiota from macroalgae were sampled each month during low tides with a high tidal coefficient (> 80) between February 2020 and January 2021. Due to the Covid-19 pandemic, no sampling was done in April and May 2020. Microbiota were collected using sterile swabs (Zymobiomics) on healthy adult specimens of the brown algae *Laminaria digitata* (Ldig, blade length 0.5-1 m), *Fucus serratus* (Fser) and *Ascophyllum nodosum* (Anod), the red alga *Palmaria palmata* (Ppal) and the green alga *Ulva* sp. (Ulva) at the Bloscon site (48°43’29.982’’ N, 03°58’8.27’’ W) in Roscoff (Brittany, France). Three individuals of each species were sampled. Additionally, samples from stranded algae (Std) were retrieved: within the mixture of diverse decomposing algae, microbiota from one *L. digitata*, one *F. serratus* and one *P. palmata* specimens were collected each month. Swabbed surface was standardized to 50 cm^2^ using a 5-cm square template on both sides of the algal thallus when possible. For *Ascophyllum nodosum*, a 1 cm-width frond was sampled on 25 cm on both sides. Three different blade areas of the kelp *L. digitata* were sampled: the basal part (young tissue, hereafter LdigB and "base"), the medium frond (ca. 20 cm away from the stipe/blade junction, hereafter LdigM and "medium") and the old frond (the blade tip, hereafter LdigO and "old"). The different algal surfaces sampled (different species and different thallus parts) are referred to as “compartments” in the analyses. Swabs were immersed in DNA/RNA Shield reagent (ZymoBiomics), kept on ice during transport and stored at -20 °C until DNA and RNA extraction.

Natural tidally-mixed coastal surface seawater (1 m depth) was collected every two weeks, during high neap tides (tidal coefficient < 60), from 2009 to 2016 at the SOMLIT-Astan station in the western English Channel, 3.5 km off Roscoff (France, 48°46′40″ N, 3°56′15″ W) along with samples collected in the frame of the SOMLIT monitoring program (Service d’Observation en Milieu Littoral; http://www.somlit.fr). Seawater was collected using 5-liters Niskin bottles and transported to the laboratory in 10-liters Nalgene bottles. Seawater samples (5 l) were filtered onto successive 3 µm and 0.2 µm polycarbonate membranes (47 mm, Whatman) that were preserved in 1.5 ml of lysis buffer (sucrose 256 g.l^-1^, Tris-HCl 50 mM pH 8, EDTA 40 mM) and stored at -80°C until further processing (Caracciolo *et al*., 2022).

### Nucleic acids extraction

Environmental DNA and RNA from swabs on macroalgae were extracted simultaneously using the DNA/RNA Miniprep kit (ZymoBiomics). All samples were processed between December 2020/January 2021, except samples from February 2020 that were processed two weeks after collection. Briefly, samples were transferred from DNA/RNA Shield – Lysis Tube to a 2 ml eppendorf and cells lysed using a TissueLyser II (Qiagen) at maximum speed (5 repetitions of 1 min with 1 min on ice between each). Tubes were centrifuged (16000 *g*, 1 min), supernatant transferred in a new tube and one volume of lysis buffer was added.

Samples were transferred on the provided Spin-Away filter and centrifuged. DNA binds to the filter while RNA is in the flow-through. The latter was collected, mixed with 1 volume of 96 % molecular biology grade ethanol, loaded on the provided Zymo-Spin IIICG column and centrifuged. RNA was treated with DNase I Reaction Mix for 15 min at room temperature. DNA and RNA were purified simultaneously, the column matrix was washed twice and nucleic acids eluted in 50 μl RNase-free water. DNA and RNA were stored at -20 and -80 °C, respectively.

For coastal seawater samples, DNA was extracted from filters as described previously (Caracciolo *et al*., 2022) with cell lysis for 45 min at 37 °C using 100 µl lysozyme (20 mg.ml^-1^) and 1 h at 56 °C with 20 µl proteinase K (20 mg.ml^-1^) and 100 µl SDS 20 %, followed purification by phenol-chloroform methods and silica membranes from the NucleoSpin PlantII kit (Macherey-Nagel).

DNA concentration was assessed with the QuantiFluor dsDNA System (Promega) kit and samples were normalized at 0.5 ng.μl^-1^ before qPCR assays. RNA concentration was measured using a Qubit instrument (Thermofisher) with the RNA HS Assay Kit and samples were normalized at 1 ng.μl^-1^ before reverse transcription.

### qPCR and RT-qPCR assays

PCRs were carried out in 384-multiwell plates on a LightCycler 480 Instrument II (Roche) as described in Brunet et al., 2021, using either universal bacterial primers 926F/1062R (Klindworth *et al*., 2013) or *Zobellia*-specific primers 142F/289R ( Brunet *et al*., 2021) to assess total and *Zobellia* 16S rRNA gene copies, respectively. Each 5 μl reaction contained 2.5 μl of LightCycler 480 SYBR Green I Master mix 2X (Roche Applied Science), 0.5 μl of each primer (300 nM final) and 1.5 μl of template DNA. Each reaction was prepared in technical triplicates. The amplification program consisted of an initial denaturation at 95 °C for 10 min followed by 45 cycles of 95 °C for 10 s, 20 s at the chosen annealing temperature (60 and 64 °C for the universal and the *Zobellia*-specific primers respectively), and polymerization at 72 °C for 10 s. After the amplification step, dissociation curves were generated by increasing the temperature from 65 °C to 97 °C. A dilution series of purified *Z. galactanivorans* Dsij^T^ gDNA (prepared as in Gobet *et al*., 2018) representing from 10 to 10^8^ 16S rRNA gene copies was used as a standard curve and were amplified in triplicate in the same run as the environmental samples. Non-Template Controls (NTC) were included in the run. The LightCycler 480 Software v1.5 (Roche) was used to determine the threshold cycle (Ct) for each sample. Linear standard curves were obtained by plotting Ct as a function of the logarithm of the initial number of 16S rRNA gene copies. PCR efficiency was calculated as 10^-1/slope^ - 1. Samples with Ct outside the linear portion of the curve were considered as null. RT-qPCR was carried out for samples from *L. digitata* and from stranded alga. Before RT-qPCR, PCR reactions were carried out with the universal primers to ensure the absence of DNA contamination. Reverse transcription was conducted on the normalized RNA samples using the ImProm-II Reverse Transcription System (Promega). After addition of 1 μl of random primers to 9.5 μl of sample, RNA was denaturated and reverse transcripted by adding 10.5 μl of the reaction mix (4 μl of 5X Reaction Buffer, 3 μl of 25 mM MgCl_2_, 1 μl of 10 mM (of each dNTP) dNTP Mix, 0.5 μl of 40 U.μl^-1^ Ribonuclease Inhibitor and 1 μl of Reverse Transcriptase) and incubating the mix for 5 min at 25 °C, followed by 1 h at 42 °C and 15 min at 70 °C. qPCR was conducted immediately after reverse transcription with 1 μl of 10X diluted cDNA as described above (0.5 μl of molecular-grade water was added per reaction to have a final volume of 5 μl). MIQE information related to (RT-)qPCR experiments is available in SuppTable 1.

### Hydrological data from the water column

All hydrological data used in this study were retrieved from the SOMLIT database (Service d’Observation en Milieu Littoral; http://www.somlit.fr, Cocquempot *et al*., 2019) on April 7th, 2022, from the SOMLIT-Astan station (period 2009-2016) and the SOMLIT-Estacade station (period 2020-2021, 1.3 km from the macroalgae sampling site, 48°43′56″ N, 3°58′58″ W). Datasets include 15 hydrological parameters, namely surface seawater temperature (T, °C), salinity (S), dissolved oxygen (O, ml.l^-1^), pH, concentration of ammonium (NH_4_, μM), nitrate (NO_3_, μM), nitrite (NO_2_, μM), phosphate (PO_4_, μM), silicate (SiOH_4_, μM), particulate organic carbon (COP, μg.l^-1^) and nitrogen (NOP, μg.l^-1^), suspended matter (MES, mg.l^-1^), ^15^N (DN15, °/_°°_) and ^13^C isotopes (DC13, °/_°°_) and Chlorophyll a (CHLA, μg.l^-1^). Protocols are available on the SOMLIT website and described in Gac *et al*. (2020).

### rRNA metabarcoding data for protists in the water column

The Operational Taxonomic Units (OTU) table of publicly available 18S V4 rRNA metabarcoding data for protists in the size fraction > 3 µm at the SOMLIT-Astan station (period 2009-2016) was retrieved on December 9th 2022 (https://zenodo.org/record/5032451#.Y5Maw-zMJTY). Pipelines for the generation of OTUs from raw reads and the OTU table were described previously (Caracciolo *et al*., 2022) and available at https://doi.org/10.5281/zenodo.5791089. The dataset comprised 21,418 protist OTUs with over 30 million sequence reads.

### Statistical analysis

For each sample, the number of *Zobellia* 16S rRNA gene copies was divided by that obtained with universal bacterial primers to assess the relative abundance of *Zobellia*. Statistical analyses were performed with R v3.5.0 (R Core Team, 2018) with significance threshold α = 0.05. For macroalgal surface samples, 2-way ANOVA with interaction effect were performed to test the effect of macroalgal compartments and sampling months on the abundance of *Zobellia*, followed by pairwise post-hoc Tukey HSD when significant results were found. Additionally, one-way ANOVA analyses were conducted to test the effect of macroalgal status ("healthy" vs. "stranded") on the abundance of *Zobellia*. Pearson correlation coefficients were calculated between averaged triplicates of *Zobellia* absolute and relative abundance on macroalgal surfaces and the different parameters using the corrplot package (v0.90). Correlations were considered significant when Benjamini-Hochberg adjusted p-value was < 0.05.

For water column samples, the difference between the abundance of *Zobellia* in free-living vs. particle-associated fractions was tested using the Mann-Whitney test. The seasonal effect was tested using Kruskal-Wallis tests followed by post-hoc Dunn test with Benjamini-Hochberg corrections of P-values. Time-dependent associations between the relative abundance of *Zobellia* in the fraction > 3 µm and hydrological parameters or relative abundance of eukaryotic OTUs were searched using extended Local Similarity Analysis (eLSA v1.0.2) (Xia *et al*., 2011). A prevalence filter of 50 % was applied to keep only eukaryotic OTUs found in at least half of the 188 samples for analysis (Röttjers and Faust, 2018). eLSA was run with a maximum delay of 3 successive sampling events, 1,000 permutations, the "nearest" method to fill missing values and "percentileZ" normalization. Local Similarity scores were standardized on a [-1;1] scale (Xia *et al*., 2011) and are referred to as stdLS. Associations were considered significant when covering more than 75 % of the time series (length > 141 successive sampling events) and |stdLS| ≥ 0.6 and P<0.001 and Q<0.05.

## Results

### Seasonal variations of *Zobellia* abundance and activity on different macroalgal surfaces

Quantitative PCR was carried out for the 238 DNA samples from algal microbiota to assess the seasonal prevalence and abundance of the genus *Zobellia* on healthy and stranded macroalgae with distinct chemical composition (SuppTable 2, Figure 1). The sample LdigB_1 in June 2020 displayed 52,100 *Zobellia* 16S rRNA gene copies.cm^-2^, which is 3.2 times more than in the sample with the second highest abundance (16,300 *Zobellia* copies.cm^-2^). After boxplot analysis of all results, it was considered an outlier and was then removed from further analyses. Two-way ANOVA revealed no interaction effects between the sampled compartment and collection month for the number of total copies and the proportion of *Zobellia* copies (F_36,158_ = 0.7, P = 0.9 and F_36,158_ = 1.1, P = 0.4, respectively) while a slight effect was detected for the number of *Zobellia* copies (F_36,158_ = 2.8, P < 0.001).

**Figure 1:**
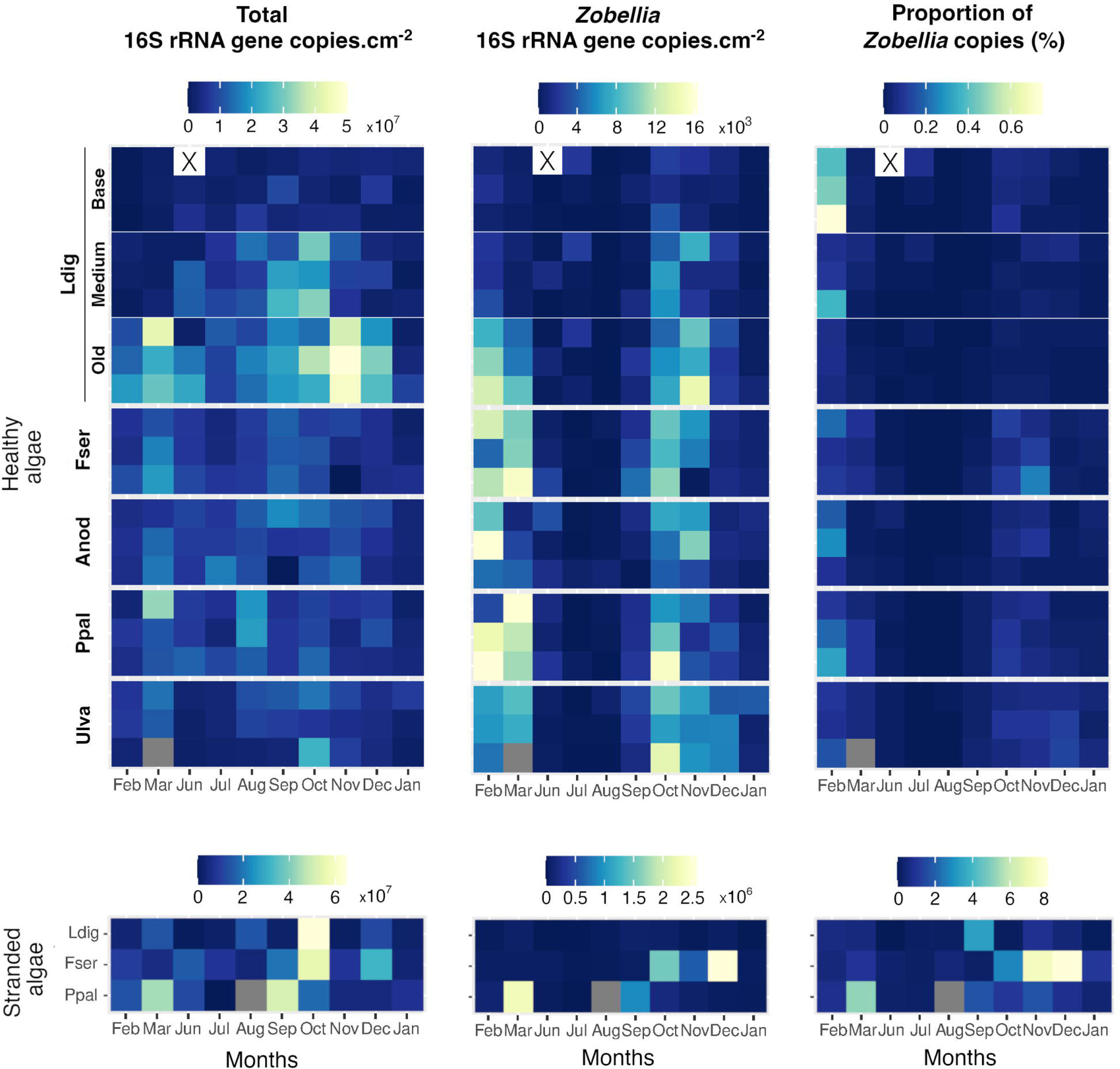
Abundance of total and *Zobellia* 16S rRNA gene copies at the surface of healthy (top) and stranded (bottom) macroalgae. The number of total (left) and *Zobellia* (middle) 16S rRNA gene copies per cm^2^ is shown, as well as the proportion of *Zobellia* copies (right). No data are available for *Ulva*_3 in March as the sample was lost. The sample LdigB_1 is considered as an outlier as 52,100 *Zobellia* copies were measured, and it was not considered for statistical analyses.

16S rRNA genes from *Zobellia* were detected on the surface of the five tested healthy macroalgae, representing on average between 2,000 and 4,500 copies.cm^-2^, depending on the algal species (Figure 2A). *Laminaria digitata* harbored on average less copies of 16S rRNA genes from *Zobellia* compared to *F. serratus* (P < 0.001), *P. palmata* (P < 0.001) and *Ulva sp.* (P = 0.006). On the other hand, the relative proportion of *Zobellia* copies did not statistically differ among algae and *Zobellia* copies represented on average 0.05% of the total number of copies (Figure 2B).

**Figure 2:**
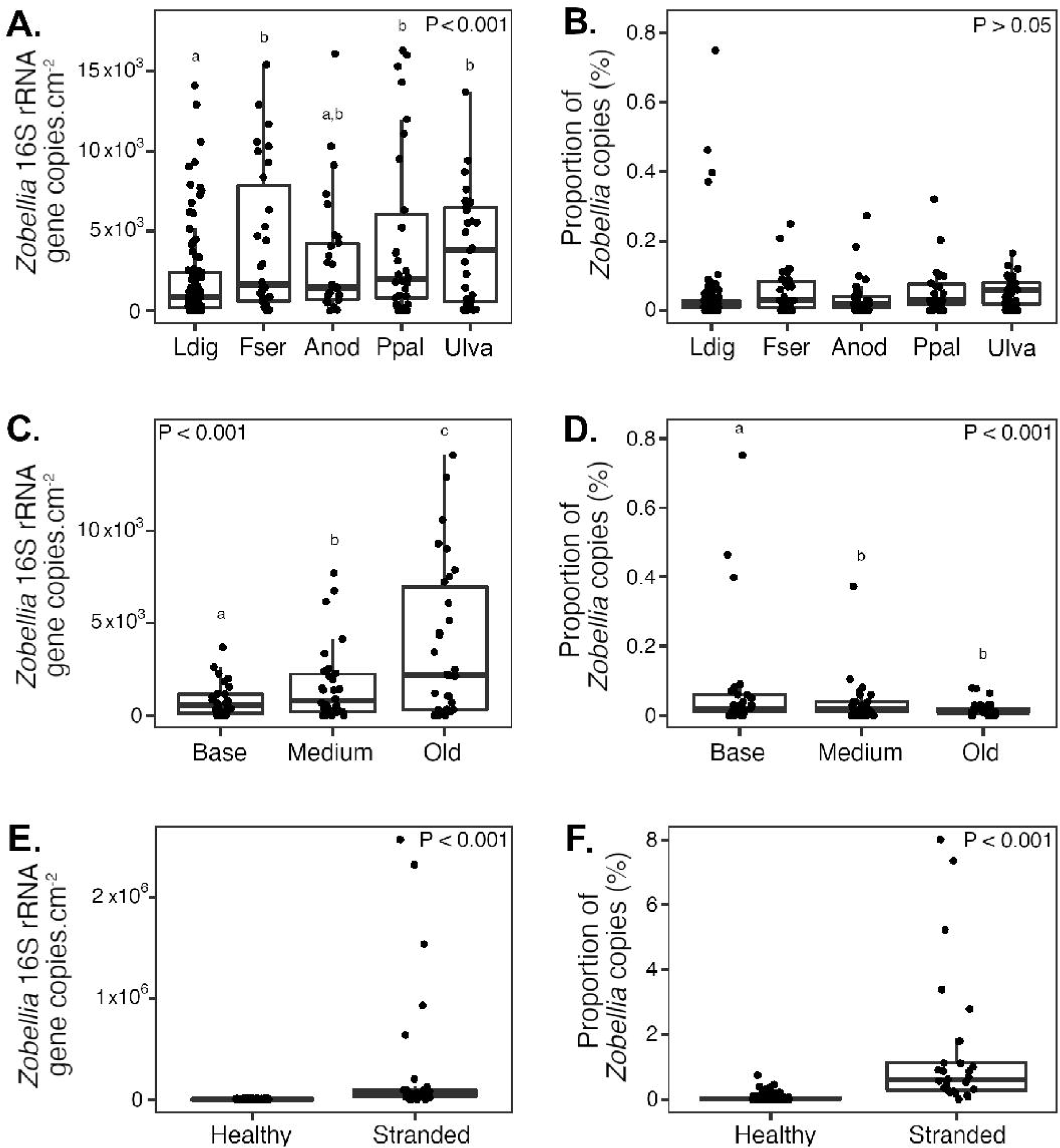
Absolute abundance (A, C, E) and relative proportion (B, D, F) of *Zobellia* 16S rRNA gene copies per cm^2^ depending on algal species (A, B), *Laminaria* blade parts (C, D) and algal status (E, F). P-values results of two-way ANOVA test followed by post-hoc Tukey HSD test are displayed, with different letters denoting significant difference (P < 0.05) across species, blade part or status.

*Zobellia* was quantified on different parts of the thallus of *L. digitata* (Figure 2C and D), revealing important variations for both the absolute amount of 16S rRNA gene copies (F_2,59_ = 43.3, P < 0.001) and their relative proportion (F_2,59_ = 12.3, P < 0.001). *Zobellia* abundance increased from the base to the tip and represented up to 10^4^ copies.cm^-2^ at the extremity of the blade - characteristic of old tissues. The proportion of *Zobellia* followed the opposite pattern as a significantly higher proportion was detected on the base (up to 0.7%) - where the tissues are renewed - than on the medium and old blade.

Analyses were also performed on stranded individuals in decomposition on the shore line (Figure 2E and F). The absolute and relative amounts of *Zobellia* 16S rRNA gene copies were highly enriched on the surface of stranded algae compared to the healthy ones (F_1,158_ = 40.2, P < 0.001 and F_1,158_ = 89.6, P < 0.001, respectively). *Zobellia* accounted on average for 1.5 % of the total number of 16S rRNA gene copies on stranded algae (30-fold more than the average on healthy tissues) and more than 5 % for three samples (*F. serratus* in November and December, *P. palmata* in March).

Contrary to the whole bacterial community, a seasonal pattern was identified for the absolute and relative abundance of *Zobellia* 16S rRNA gene copies (Figure 3, SuppFig. 1). Pairwise comparisons revealed a lower abundance of *Zobellia* during summer, from June to September. The lowest number of *Zobellia* 16S rRNA gene copies was obtained in August for *L. digitata* (ca. 0-20 copies.cm^-2^ representing 0-0.0005 % of the total copies) and in July for the other species (ca. 0-600 copies.cm^-2^ representing 0-0.002 % of the total copies). The abundance of *Zobellia* peaked twice, in February-March and October-November, with ca. 10^3^-10^4^ *Zobellia* 16S rRNA gene copies.cm^-2^, representing 0.03-0.5 % of the total amount.

**Figure 3:**
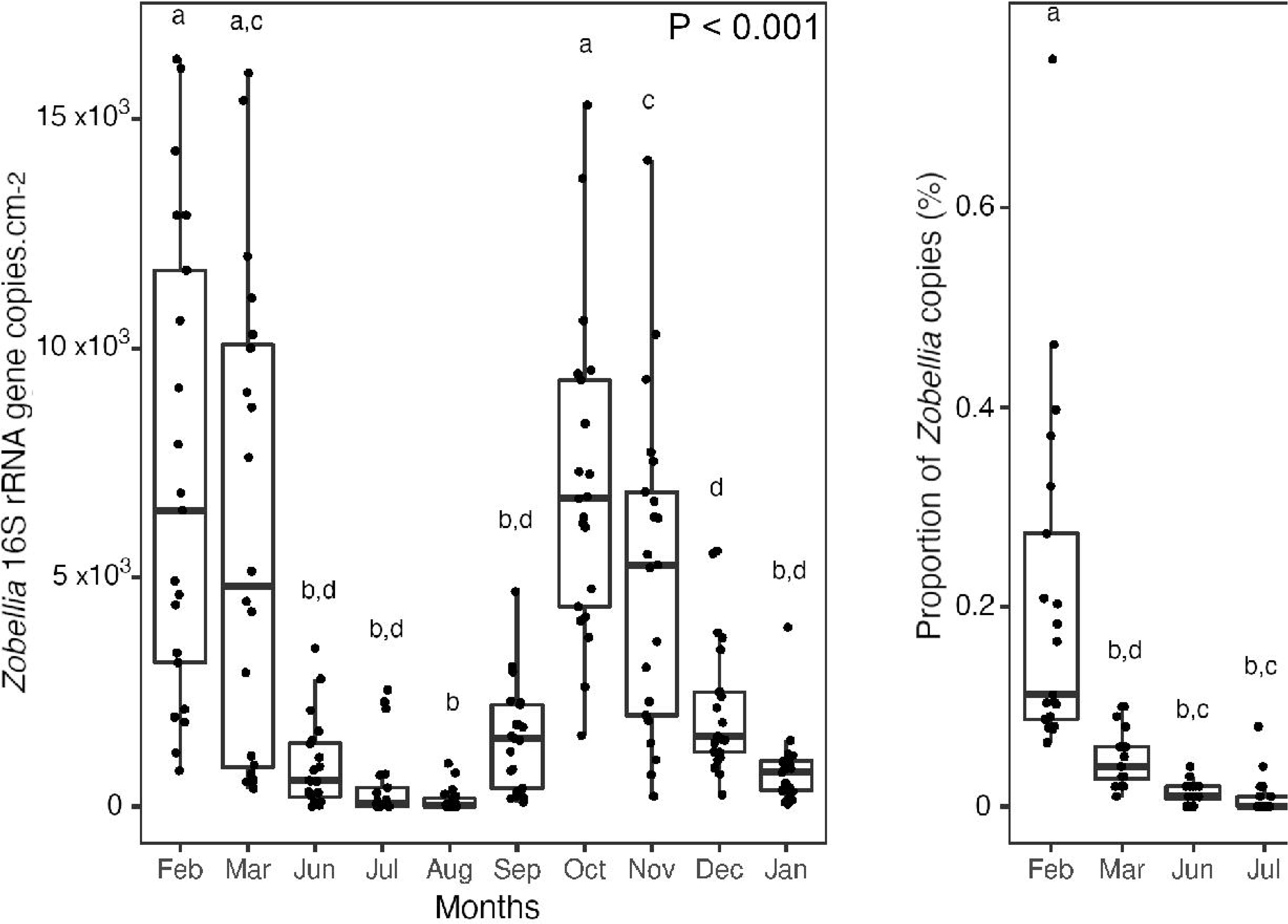
Number of *Zobellia* 16S rRNA gene copies per cm^2^ (left) and proportion of *Zobellia* copies (right) detected at different sampling months between February 2020 and January 2021 on macroalgal tissues. P-values results of two-way ANOVA test followed by post-hoc Tukey HSD test are displayed, with different letters denoting significant difference (P < 0.05) across months.

Among the 15 measured environmental parameters, the proportion of *Zobellia* 16S rRNA gene copies showed strongest positive correlation with concentrations of NO_3_, PO_4_, SiOH_4_ and MES and strongest negative correlation seawater temperature (T), salinity (S), pH, and concentrations of COP, NOP and Chl a, independently of the studied algae (Pearson correlation, Figure 4). Similar correlation patterns were observed for the absolute number of *Zobellia* copies (SuppFig. 2)

**Figure 4:**
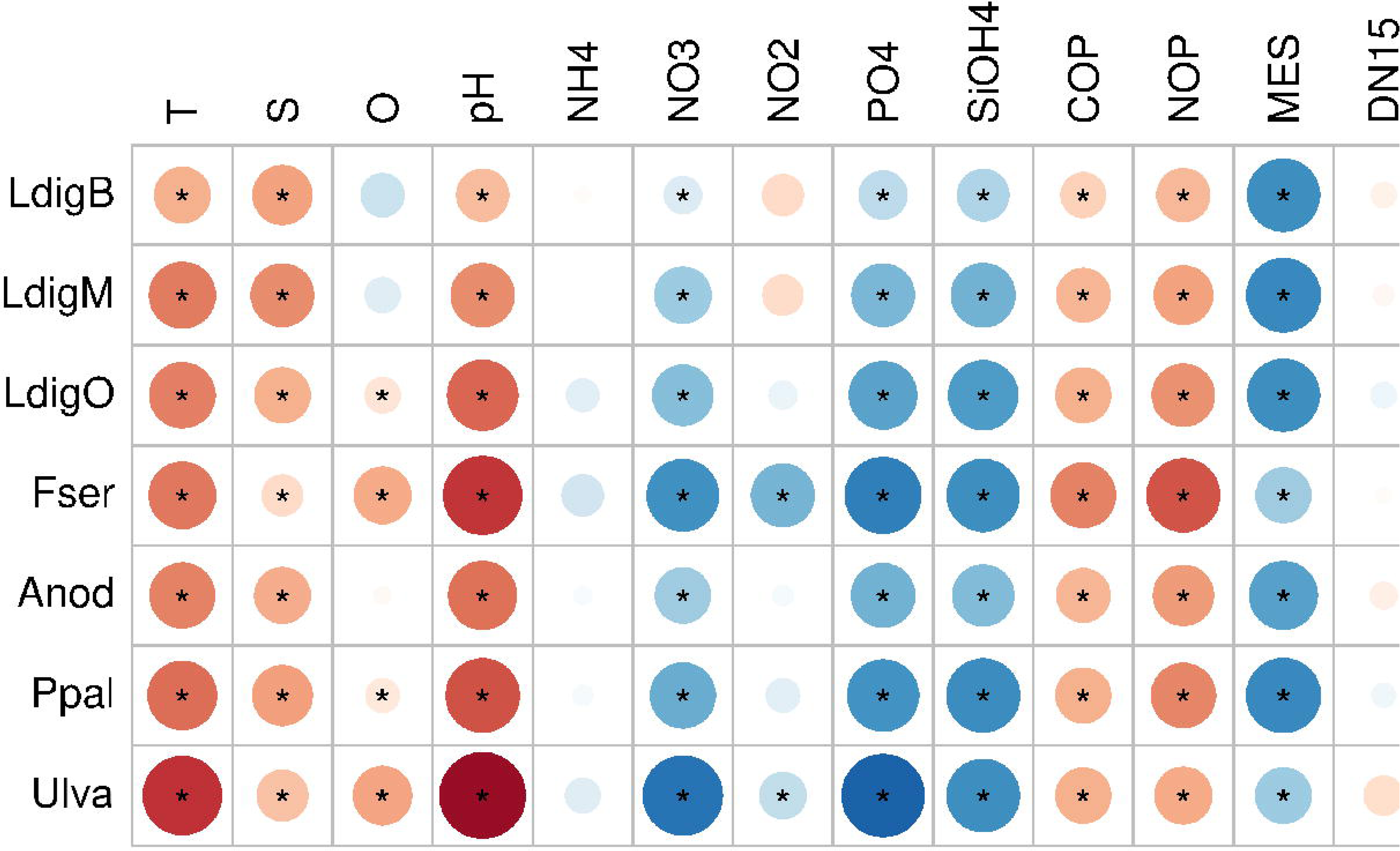
Correlation matrix (Pearson coefficients) between the proportion of 16S rRNA *Zobellia* copies on the different algal tissues and environmental parameter measurements. Asterisks denote significant correlations (Benjamini-Hochberg adjusted p-values < 0.05)

The number of 16S rRNA transcripts from total bacteria and *Zobellia* cells were estimated using RT-qPCR for *L. digitata* and stranded macroalgal samples (SuppTable 2). The ratio of 16S rRNA transcript copies over 16S rRNA gene copies (16S rRNA:rDNA ratio) was calculated as an index of activity (Campbell *et al*., 2011). When *Zobellia* 16S rRNA transcripts were detected, this *Zobellia*-specific 16S rRNA:rDNA ratio was significantly higher on stranded algae (123 ± 30, mean ± s.e.m.) compared to healthy *L. digitata* tissues (27 ± 3) (Wilcoxon test, W = 281, P < 0.001). Futhermore, the *Zobellia*-specific 16S rRNA:rDNA ratio was higher to the one calculated for the total bacterial community in the same sample on *L. digitata* tissues (2.6-fold higher, Wilcoxon paired test, W = 1558, P < 0.001), but not on stranded samples (V_26_=130, P = 0.16).

### Detection of *Zobellia* in tidally-mixed coastal waters

The presence of the genus *Zobellia* was further monitored in coastal seawater samples on a bi- weekly time series from 2009 to 2016, using qPCR both on the free-living (0.2-3 µm) and particle-associated (> 3 µm) fractions. 16S rRNA genes from *Zobellia* were detected in 35 % (50/141) and 59 % (98/166) of the free-living and particle-associated fractions, respectively (Supp Figure 3). The maximum absolute abundance of *Zobellia* was respectively 5,890 and 17,629 16S rRNA gene copies.l^-1^ in the free-living fraction (January 2011) and on particles (March 2009). Overall, *Zobellia* was 4-fold more abundant on particles (1,608 ± 257 copies.l^-1^, mean ± s.e.m, n = 166) than on the free-living fraction (473 ± 94 copies.l^-1^, mean ± s.e.m, n = 141) (Mann-Whitney test, W = 8,302, P < 0.001). The same trend was observed for the relative abundance of *Zobellia* within the total bacterial community (Mann-Whitney test W = 7,783, P < 0.001). On particles, *Zobellia* accounted for up to 0.21% of the total bacterial community. Both the absolute and relative abundance of *Zobellia* showed a recurring seasonal trend, with significantly higher values during the winter (Figure 5). To explain this seasonal pattern, we used extended Local Similarity Analysis (eLSA) to detect associations between the relative abundance of *Zobellia* and hydrological parameters, focusing only on the > 3 µm particles were the presence of *Zobellia* was more consistent. The strongest positive and negative associations were found for nitrate concentration (stdLS = 0.952) and seawater temperature (stdLS = -1), respectively (Figure 6A). In addition, the relative abundance of *Zobellia* was positively associated to silicate, oxygen and phosphate concentrations, while it was negatively associated to salinity, chlorophyll a and ammonium concentrations (SuppTable 3). eLSA also revealed positive associations between the relative abundance of *Zobellia* and that of 114 individual eukaryotic OTUs (SuppTable 3). Most positive associations were found with Stramenopiles (37 OTUs, 32 %) and Alveolata (35 OTUs, 31 %). The relative abundance of *Zobellia* was notably associated positively without delay to 3 brown macroalgal OTUs within the Phaeophyceae and one red macroalgal Corallinales OTU (Figure 6B). Furthermore, strong positive associations were detected with protist OTUs (Figure 6C), including Ventrifissuridae (highest stdLS = 1), Parmales (highest stdLS = 0.951), Dinophyceae (highest stdLS = 0.945), *Chloropicon* (highest stdLS = 0.921) and Syndiniales (highest stdLS = 0.897).

**Figure 5:**
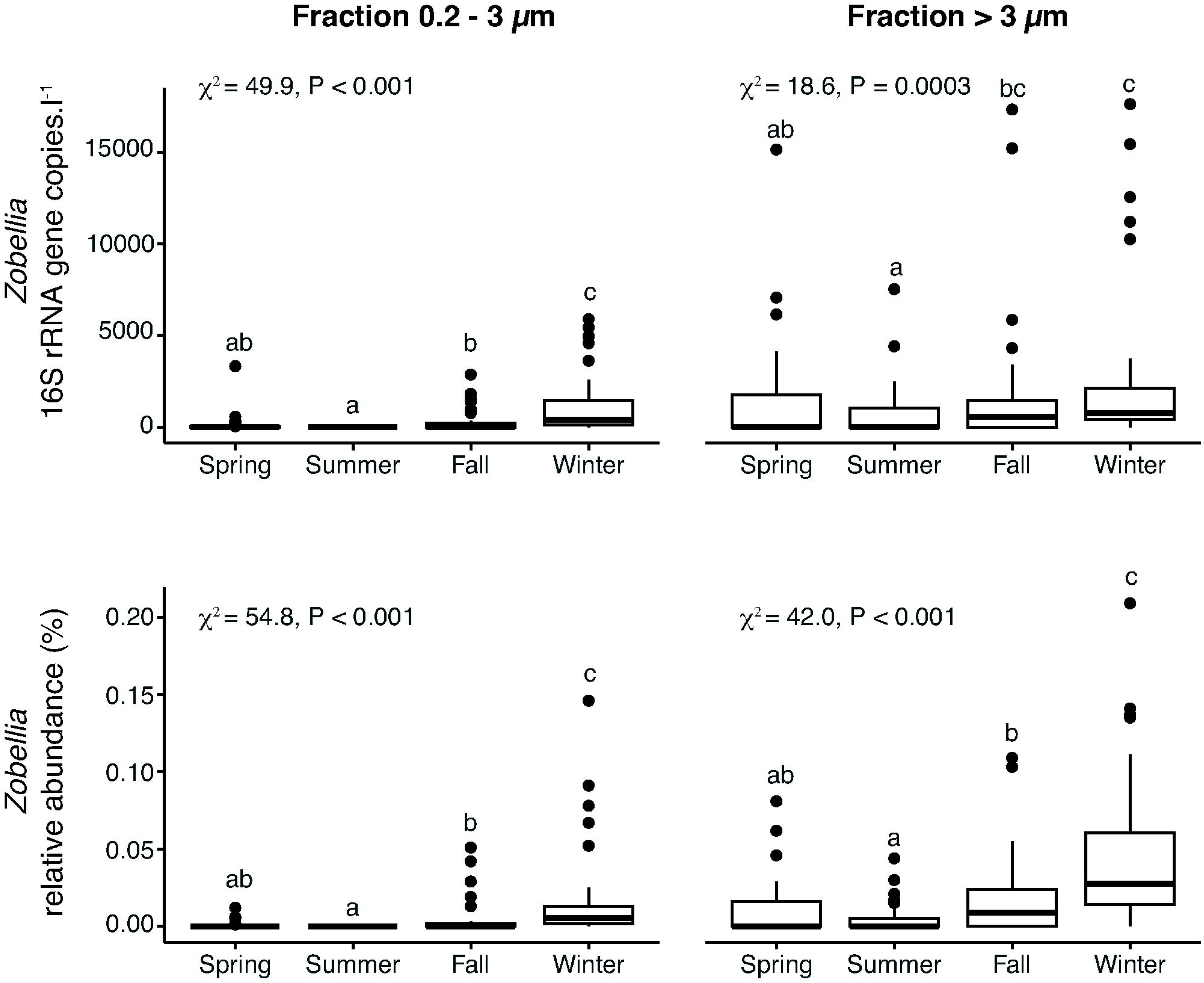
Seasonal effect on the absolute (top row) and relative (bottom row) abundance of *Zobellia* in free-living (0.2 – 3 µm, left lane) and particle-associated (> 3 µm, right lane) fractions. Seasons were defined as follows: Spring, ordinal days 81-172; Summer, ordinal days 173-264; Fall, ordinal days 265-355; Winter, ordinal days < 81 and ≥ 356. χ^2^ and P-values results of Kruskal-Wallis test followed by post-hoc Dunn test of the season effect are displayed, with different letters denoting significant difference between seasons.

**Figure 6:**
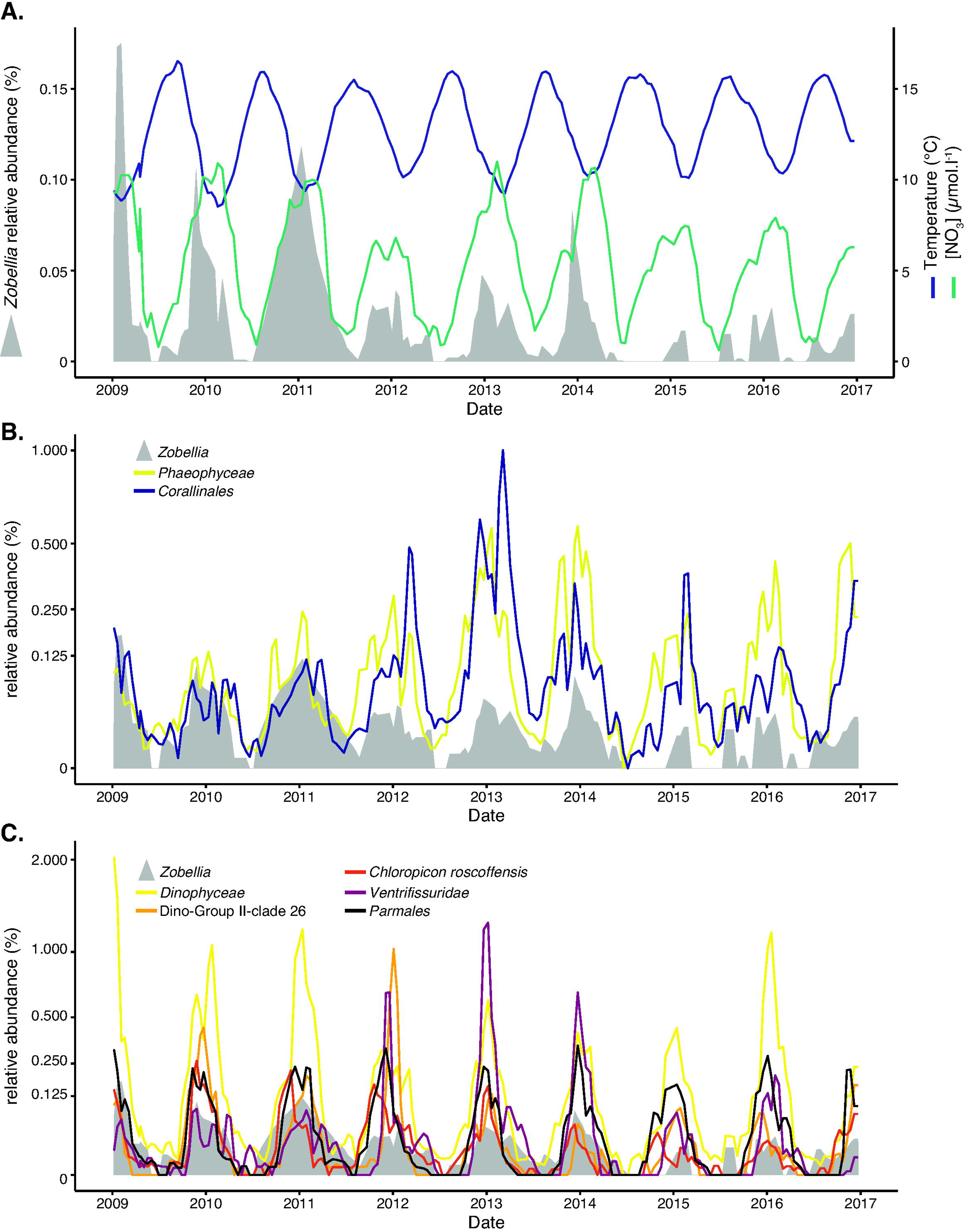
Selected significant associations of *Zobellia* with physico-chemical parameters and eukaryotic taxa in the >3 µm fraction. **A.** Relative abundance of *Zobellia* (left axis, grey area), seawater temperature (standardized LS = -1, length 185, delay = -2, right axis) and nitrate concentration (std LS = 0.952, length = 176, delay = -1, right axis). **B.** Relative abundance of *Zobellia* (grey area) and two selected macroalgal OTUs with significant positive association, affiliated respectively to *Phaeophyceae* (std LS = 0.687, length 187, delay = 0) and *Corallinales* (std LS = 0.681, length = 187, delay = 0). **C.** Relative abundance of *Zobellia* (grey area) and five selected protist OTUs with significant positive association, affiliated respectively to *Dinophyceae* (std LS = 0.945, length = 187, delay = 0), Dino Group II – clade 26 (std LS = 0.897, length = 187, delay = 0), *Chloropicon roscoffensis* (std LS = 0.921, length = 187, delay = 1), *Ventrifissuridae* (std LS = 1, length = 182, delay = 0) and *Parmales* (std LS = 0.951, length = 187, delay = 1). Values are rolling means (window width = 2 consecutive sampling events). For each year, the tick mark corresponds to January 1^st^. Full results are available in SuppTable 3.

## Discussion

### 1) *Zobellia* is a stable component of healthy macroalgal epibiota

Our study shows that the flavobacterial genus *Zobellia* is consistently part of the epibiota of macroalgae in a temperate coastal habitat. This agrees with the fact that most of the described *Zobellia* species have been isolated from macroalgae (Barbeyron *et al*., 2001, 2021; Nedashkovskaya *et al*., 2004; Nedashkovskaya *et al*., 2021) and that *Zobellia* sequences were found in metagenomic studies on macroalgae (Miranda *et al*., 2013; Dogs *et al*., 2017). Here, *Zobellia* was detected year-round on the five algal species tested, which cover all three macroalgal phyla and whose chemical composition largely differs between one another (Kloareg *et al*., 2021). Given its stable presence on the diverse algal species, *Zobellia* might therefore display neutral or beneficial interactions with healthy hosts. Indeed, *Zobellia* representatives are known to possess multiple adaptive traits that may favor stable association with macroalgal hosts (Barbeyron *et al*., 2016). It includes resistance mechanisms against oxidative stress and halogenated compounds (Fournier *et al*., 2014; Grigorian *et al*., 2021) and the secretion of thallusin, a macroalgal morphogenetic compound (Ulrich *et al*., 2022). Furthermore, described *Zobellia* species can assimilate several of the main glycans produced by brown (eg. alginate, fucan-containing sulfated polysaccharides, laminarin, mannitol), red (eg. agar, porphyran, carrageenans) and green macroalgae (eg. ulvans) (Barbeyron *et al*., 2001, 2021; Nedashkovskaya *et al*., 2004; Groisillier *et al*., 2015). This metabolic versatility might be an advantage to colonize diverse macroalgal tissues offering different substrate niches.

Despite this stable association with healthy macroalgae, *Zobellia* was not a dominant genus compared to other taxa such as *Granulosicoccus* spp. (*Gammaproteobacteria, Chromatiales*) or *Litorimonas* spp. and *Hellea* spp. (*Alphaproteobacteria, Maricaulales*) that can represent more than 10 % of the biofilm community on kelps (Paix *et al*., 2019; Ramirez-Puebla *et al*., 2020; Brunet *et al*., 2021). Here, we show that *Zobellia* accounted on average only for 0.05% of the total bacterial community, but reached up to 0.7% depending on the algal species and season (see below). Although small, this proportion appears ecologically relevant regarding the hundreds of different taxa reported to live on macroalgal surfaces (Egan *et al*., 2013). 16S rRNA:rDNA ratios also suggest *Zobellia* cells might be more transcriptionally active than the average bacterial epibionts on healthy macroalgae. In addition, living macroalgae might actively control the development of such potent degraders as *Zobellia*, known for their large CAZyme repertoires (Barbeyron *et al*., 2016; Chernysheva *et al*., 2019) and efficiency to utilize algal biomass (Brunet *et al*., 2022).

The abundance of *Zobellia* also varied along the thallus of the kelp *L. digitata*. As observed for the total bacterial community, the absolute amount of *Zobellia* 16S rRNA genes increased with the distance from the stipe/blade junction, being maximum at the distal ends. Such a pattern in bacterial abundance was already reported for several kelp species (Staufenberger *et al*., 2008; Weigel and Pfister, 2019; Ihua *et al*., 2020). Kelps grow from the meristematic region between the perennial stipe and the blade. Therefore, the base of the blade represents the youngest tissues that might not yet host as many bacterial cells as older tissues towards the apex. On the other hand, physical shedding of distal blades likely eases the access to algal compounds, favoring bacterial growth. By contrast, the relative abundance of *Zobellia* followed an inverse trend, being significantly higher on the basal blade of *L. digitata*. Previous studies have shown that younger tissues of several kelp species release more reactive oxygen species upon stress (Küpper *et al*., 2001) and contain higher concentrations of phlorotannins (Van Alstyne *et al*., 1999), which can both control the growth of epiphytic bacteria. Our results therefore suggest that *Zobellia* cells might better tolerate these algal defense reactions than other bacterial epibionts on young kelp tissues.

### 2) The abundance and activity of *Zobellia* increase on stranded algae

Throughout the year, both the absolute and relative abundance of *Zobellia* were higher on stranded macroalgae compared to healthy ones. This suggests that *Zobellia* representatives become a dominant component of the bacterial community associated with decaying algal tissues, showing how competitive they can be to exploit this substrate-rich ecological niche. Together with their large genomes (ca. 5 Mb) and extended CAZyme repertoires, this illustrates their copiotrophic lifestyle, as reported for other flavobacteria (Lauro *et al*., 2009). Members of the genus *Zobellia* also feature additional traits that might partly help them compete with other members of the bacterial community, namely the synthesis of the dialkylresorcinol antimicrobial zobelliphol active against Gram-positive bacteria (Harms *et al*., 2018) and the predicted production of a homoserine lactone lyase putatively interfering with quorum sensing mechanisms (Barbeyron *et al*., 2016). We further showed that the 16S rRNA:rDNA ratio was high for *Zobellia* on stranded macroalgae, illustrating a strong activity. Since ribosome content correlates positively with growth rate for many bacteria, rRNA:rDNA ratio is a common proxy to estimate growth rates of specific bacterial taxa in natural communities (Campbell *et al*., 2011). Based on linear regression analysis of rRNA:rDNA versus growth rate for various marine and non-marine copiotrophic bacteria (Lankiewicz *et al*., 2016), we can extrapolate that the measured rRNA:rDNA ratio for *Zobellia* on stranded macroalgae would correspond to growth rates from 0.001 to 0.1 h^-1^. This matches maximum growth rates previously measured on batch cultures of *Z. galactanivorans* Dsij^T^ during the exponential phase in rich medium (Thomas *et al*., 2011). Hence, our estimate of high growth rates for natural populations of *Zobellia* on stranded macroalgae could further explain how they become dominant. We recently showed that *Zobellia* isolates actively degrade kelp tissues in laboratory microcosms (Brunet *et al*., 2022). The present data further indicates that *Zobellia* could significantly influence the fate of carbon sequestered by macroalgae in the studied coastal area, since fast-growing bacteria might inject more organic carbon into the microbial loop and produce more CO_2_ through respiration than slow growers (Del Giorgio and Cole, 1998; Pedler *et al*., 2014).

### 3) *Zobellia* is also part of coastal bacterioplankton

Although the genus *Zobellia* is hitherto mostly known for its association with macroalgal surfaces, our results show that it is also recurrently found in coastal bacterioplankton of the Western English Channel. The abundance of *Zobellia* was higher on particles than in the free-living fraction, in agreement with its ability to adhere to, glide and form dense biofilms on surfaces (Salaün *et al*., 2012; Nedashkovskaya and Suzuki, 2015). The relative abundance of *Zobellia* was in the same order of magnitude on healthy macroalgae and on particles, suggesting the coastal water column might be a yet overlooked habitat for this genus. Indeed, one of the eight validly described species, *Z. amurskyensis* KMM 3526^T^, was isolated from seawater in Amursky Bay, Sea of Japan (Nedashkovskaya *et al*., 2004). *Zobellia* representatives were also previously grown from seawater and particles larger than 10 µm during a phytoplankton spring bloom off Helgoland (Heins and Harder, 2023). Our analysis of long-term observation data revealed recurring associations of *Zobellia* with individual eukaryotic OTUs in the particle-associated fraction. This contrasts with previous studies of free-living bacterioplankton communities during coastal spring phytoplankton blooms over 4 years, which did not evidence clear correlations between distinct bacterial and diatom taxa (Teeling *et al*., 2016). Specific one-to-one interactions might therefore have a stronger influence on bacterial community composition for particle-associated clades, while different bacterial taxa sharing similar functional traits might at least partially substitute each other in the free-living fraction (Teeling *et al*., 2016; Galand *et al*., 2018). Here, we notably detected significant associations of *Zobellia* with individual OTUs affiliated to brown and red macroalgae in the particle-associated fractions. This indicates that *Zobellia* could thrive on macroalgal debris released in the water column. However, stronger associations were found with phytoplankton OTUs, including dinoflagellates, green microalgae, diatoms and Parmales (Stramenopiles, Bolidophyceae). Therefore, *Zobellia* might also be part of phytoplankton microbiomes in coastal waters, confirming previous cultivation attempts that retrieved *Zobellia* isolates from North Sea phytoplankton (Hahnke and Harder, 2013). Phytoplankton-derived polysaccharides, accounting for up to 90% of microalgal exudates (Myklestad, 1995), could be substrates for *Zobellia* growth. Recent analyses of diatom exopolysaccharides notably revealed the presence of a complex cocktail of glycans, including epitopes recognized by antibodies that target fucose-containing sulfated polysaccharide (FCSP), β-1,4-mannan, β- 1,3-glucan, xyloglucan, β-1,4-xylan, and alginate (Huang *et al*., 2021). Other detected associations of *Zobellia* with non-photosynthetic protists could indicate grazing interactions, for example for heterotrophic Stramenopiles flagellates known to graze on flavobacteria (Massana *et al*., 2009), or sharing of the same habitat, for example for heterotrophic Cercozoa that can feed on diatoms (Drebes *et al*., 1996) or Syndiniales that parasite photosynthetic dinoflagellates (Guillou *et al*., 2008).

### 4) Seasonal variations

Although always representing < 1% relative abundance, *Zobellia* showed similar seasonal variations both on macroalgal surfaces and in coastal seawater from Roscoff, with maximum and minimum abundance in fall/winter and summer, respectively. This illustrates the growth and loss dynamicity of rare microbial taxa, as previously shown for coastal marine bacterioplankton in the Bay of Biscay (Alonso-Sáez *et al*., 2015). This is consistent with previous reports of the higher winter abundance of *Bacteroidota* on the kelp *Laminaria hyperborea* from Norway (Bengtsson *et al*., 2010) and on *Fucus vesiculosus* from the Baltic Sea (Mensch *et al*., 2020). Yet, seasonal changes can vary depending on site, as already shown for *Macrocystis pyrifera* epibiota (Florez *et al*., 2019). Therefore, conclusions on *Zobellia* seasonality based on sampling in Roscoff might not be directly applicable to other locations. The highest abundance of *Zobellia* in winter contrasts with previous reports of recurring growth of planktonic *Flavobacteriia* following spring phytoplankton blooms. This highlights the importance of niche specialization within marine flavobacteria (Díez-Vives *et al*., 2019) and corroborates the hypothesis that the abundance of individual taxa can be more influenced by specific interactions rather than only based on substrate availability. Two types of parameters could explain the observed seasonality of *Zobellia*: (i) abiotic hydrological variables and (ii) fluctuations of biotic factors related to the hosts. Among abiotic factors, we notably found strong negative association of *Zobellia* with seawater temperature, which ranged from 8 °C in winter to 18 °C in summer. Temperature is well-known as a crucial driver of bacterial community structure on macroalgae (Takemura *et al*., 2014; Paix *et al*., 2021) and in coastal bacterioplankton (Teeling *et al*., 2016). Recent meta-analyses of ocean microbiome datasets showed that higher seawater temperatures universally favor slow- growing taxa (Abreu *et al*., 2022), which could lead to higher competition for resources in summer. Together with potential stochastic effects, a deterministic effect of temperature could explain the observed lower peak abundance of *Zobellia* in coastal bacterioplankton in 2012 and after 2015, when winter seawater temperature stayed > 10°C. Positive associations of *Zobellia* with nitrate and other nutrients likely reflect its copiotrophic lifestyle, thriving when resources are abundant. Furthermore, cultured *Zobellia* representatives possess a nitrate assimilation pathway and can use nitrate as a sole nitrogen source (Barbeyron *et al*., 2016). Similarly, Teeling et al (2016) found that the strongest abiotic predictors of coastal bacterioplankton were temperature, salinity, silicate and nitrate concentrations. Seasonal variations in water column samples might also result from higher hydrodynamic forces during the winter in Roscoff, when more intense tidal mixing and winds produce macroalgal debris and resuspend benthic phytoplankton from sediments (Caracciolo *et al*., 2022). In addition, fluctuations of biotic factors might partly explain the observed seasonality of *Zobellia*. Algal defense chemicals such as halogenated compounds are strong selective factors for epiphytic bacterial colonizers (Goecke *et al*., 2010). The five macroalgal species studied here display maximum and minimum iodine content in winter and summer, respectively (Ar Gall *et al*., 2004; Nitschke *et al*., 2018). *Z. galactanivorans* Dsij^T^ notably accumulates high intracellular iodine concentrations and tolerates haloacids, thanks to three vanadium-iodoperoxidases, an iodotyrosine dehalogenase and a haloacid dehalogenase that have homologs in all described *Zobellia* strains (Fournier *et al*., 2014; Barbeyron *et al*., 2016; Grigorian *et al*., 2021; Grigorian *et al*., 2023). This outstanding halogen metabolism likely constitutes a selective advantage over other less-equipped bacteria in winter, when halogen concentration is highest. The growth cycle of macroalgae might also influence their epibionts. For example, kelp blade renewal and expansion occur in spring at the meristem part. In consequence, in May, the whole blade consists of fresh tissue. Hence the low proportion of *Zobellia* in summer suggests it might not be an early colonizer of macroalgae. Moreover, algal exudation is higher in summer (Abdullah and Fredriksen, 2004) and consumers of labile exudates might dominate the biofilm (Bengtsson *et al*., 2010). By contrast, the aging tissues in autumn-winter might select for specialist bacteria able to break down the complex extracellular matrix. Although this cannot be assessed with the present data, viral lysis and protist grazing also potentially control the abundance of *Zobellia*. Both factors are known to follow seasonal dynamics (Bunse and Pinhassi, 2017) and can apply selective pressure on flavobacterial taxa (Kirchman, 2002; Massana *et al*., 2009; Bartlau *et al*., 2022).

## Conclusion

These results highlighted the seasonal dynamics and environmental constraints driving natural populations of algae-degrading bacteria. Taking the well-studied genus *Zobellia* as a test case, it connects mechanistic knowledge obtained on laboratory models to potential ecological impacts on coastal carbon cycling.

## Acknowledgments

The authors thank Manon Choulot (UMR8227, Roscoff) for her help during sample collection, Nathalie Simon (UMR7144, Roscoff) for scientific supervision of the SOMLIT monitoring, Christian Jeanthon (UMR7144, Roscoff) for useful discussions on the project, Nicolas Henry (FR2424, Roscoff) for the production and publication of eukaryotic metabarcoding datasets, Thierry Cariou (FR2424, Roscoff) for integration of hydrological parameters to SOMLIT, as well as the captains and crew of the Neomysis research ship for their help during sampling at the SOMLIT-Astan and SOMLIT-Estacade stations. This work has benefited from the facilities of the Genomer platform and from the computational resources of the ABiMS bioinformatics platform (FR 2424, CNRS-Sorbonne Université, Roscoff), which are part of the Biogenouest core facility network. This work was funded by the French Government via the National Research Agency programs ALGAVOR (ANR-18- CE02-0001-01) and IDEALG (ANR-10-BTBR-04).

## Author contributions

Author contributions following the CRediT taxonomy (https://casrai.org/credit/) are as follows: Conceptualization: FT; Formal analysis: MB, FT; Funding acquisition: FT; Investigation: MB, NLD, FRJ, SR, TB, FT; Project administration: FT; Supervision: TB, FT; Visualization: MB, FT; Writing-original draft: MB, TB, FT; Writing-review and editing: MB, NLD, FRJ, SR, TB, FT.

## Conflict of interest statement

The authors have no conflict of interest to declare.

## Supplementary Figure legends

**Supp Figure 1:**
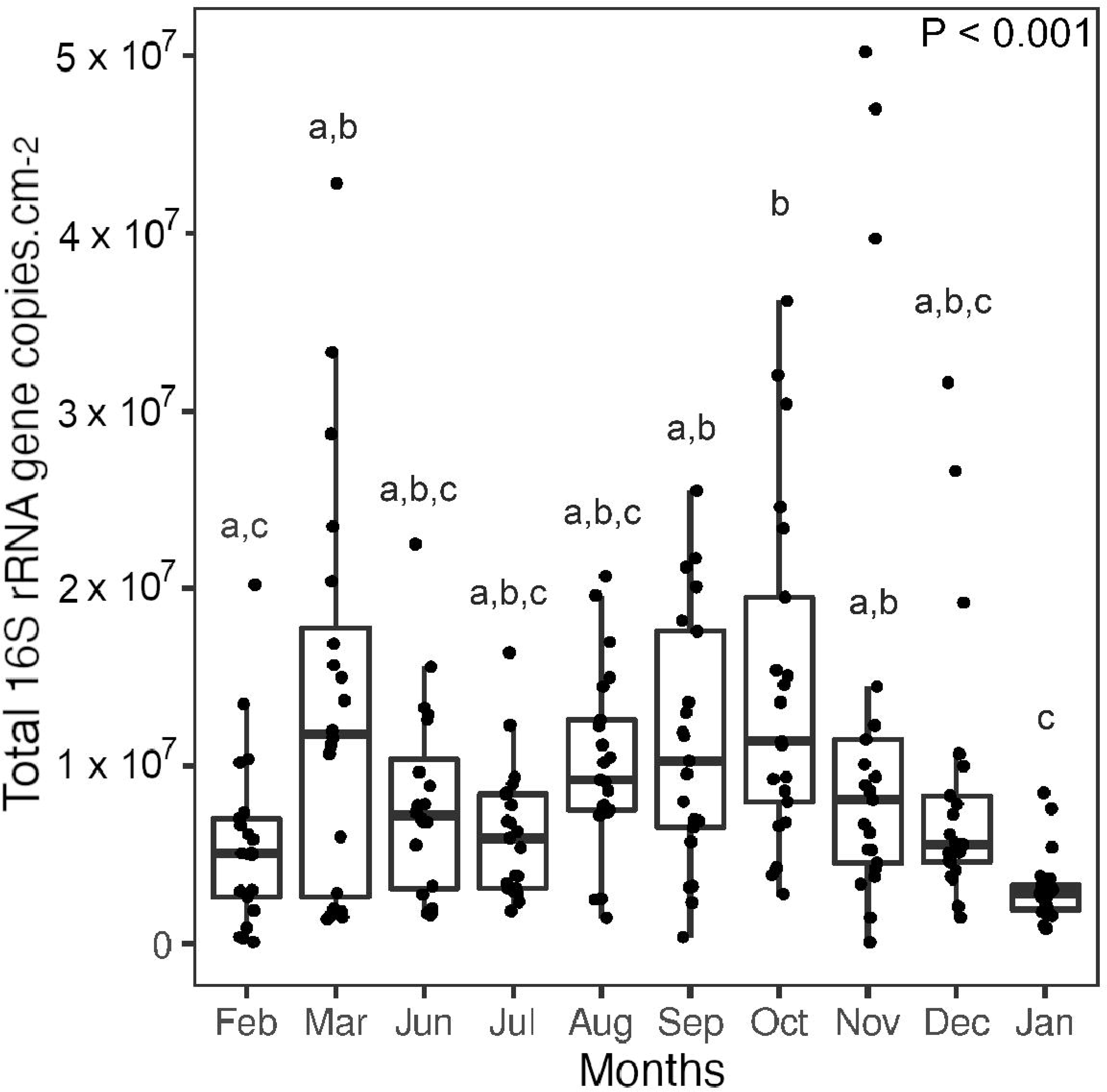
Number of total 16S rRNA gene copies per cm^2^ detected at different sampling months between February 2020 and January 2021 on macroalgal tissues. P-values results of two-way ANOVA test followed by post-hoc Tukey HSD test are displayed, with different letters denoting significant difference (P < 0.05) across months.

**Supp Figure 2:**
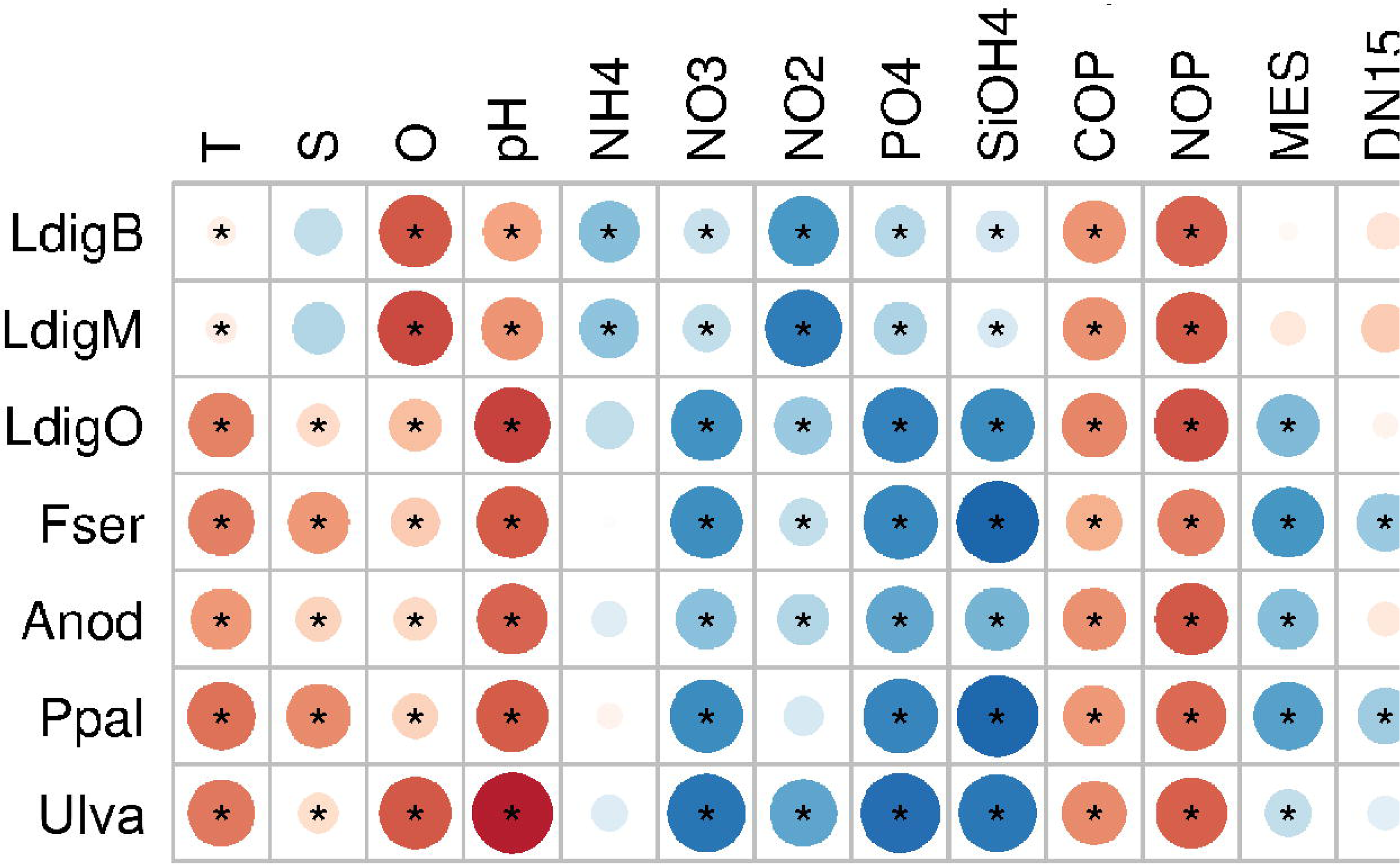
Correlation matrix (Pearson coefficients) between the number of 16S rRNA *Zobellia* copies on the different algal tissues and environmental parameter measurements. Asterisks denote significant correlations (Benjamini-Hochberg adjusted p-values < 0.05)

**Supp Figure 3:**
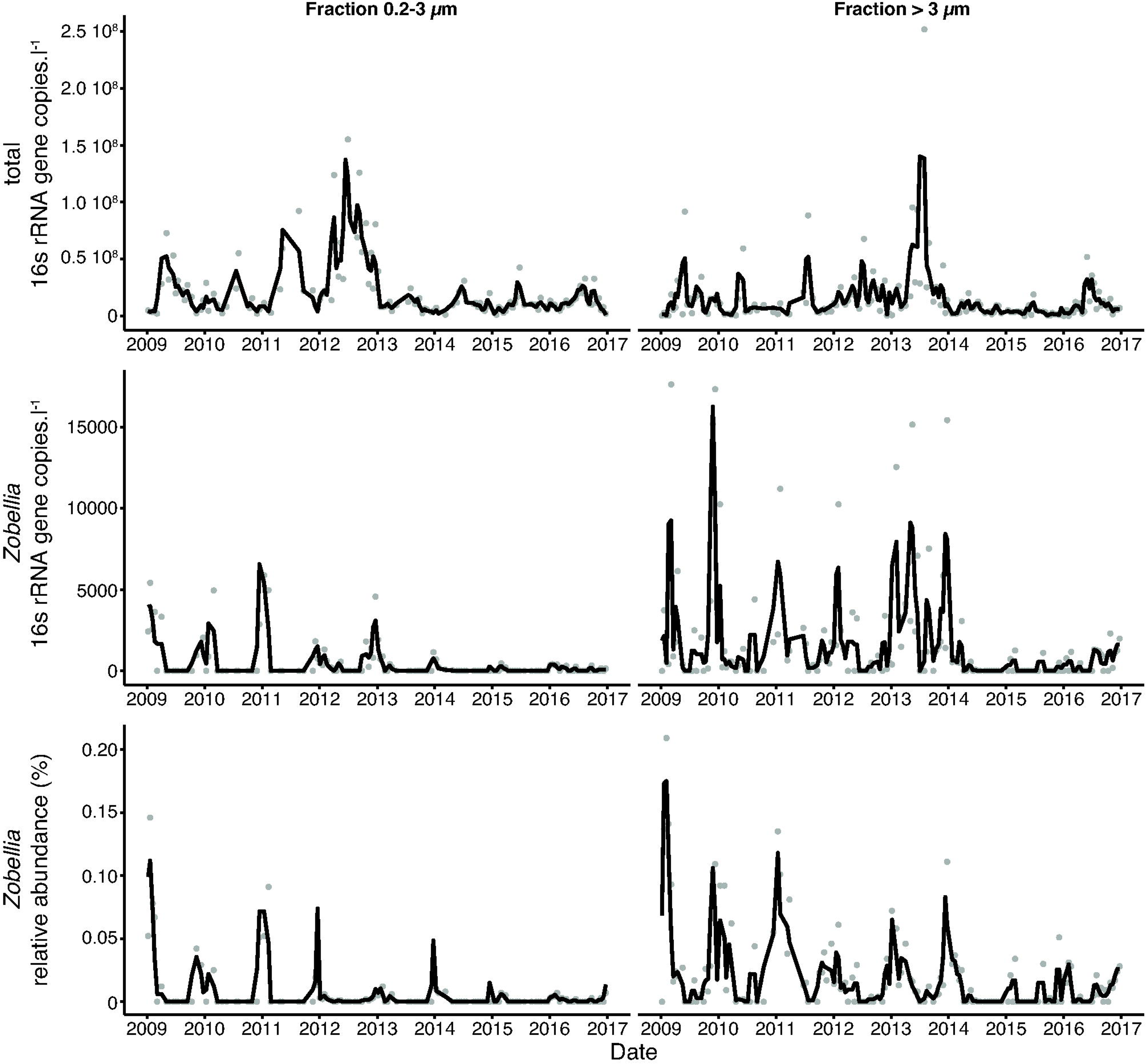
qPCR survey of the total number of 16S rRNA gene copies (top row), the *Zobellia*-specific number of 16S rRNA gene copies (middle row) and the relative abundance of *Zobellia* (bottom row) in seawater fractions below (left lane) and above 3 µm (right lane) from 2009 to 2016. For each year, the tick mark corresponds to January 1^st^. The black line represents the rolling mean (window width = 2 consecutive sampling events) and grey dots are individual measurements.

## Supplementary Tables

**Supp Table 1:** (Excel file) MIQE

**Supp Table 2:** (Excel file) Characteristics of the 238 samples of macroalgal epibiota collected during this study.

**Supp Table 3:** (Excel file) eLSA results showing significant associations of *Zobellia* relative abundance in the particle-associated fraction at the Roscoff Astan Station with hydrological parameters (both positive and negative) and protist OTUs (only positive). Parameters in red are illustrated in Figure 6. LS, Local Similarity score; stdLS, standardized LS ranging from -1 to 1. For each sample, individual values of each parameter are given for each sample, with nomenclature RAYYMMDD_3, where YY, MM and DD denote the sampling year, month and day, respectively. Hydrological parameters were retrieved from the SOMLIT database (http://somlit.fr) and protist abundances from the public dataset available at https://doi.org/10.5281/zenodo.5791089.

## Notes

### Competing Interest Statement

The authors have declared no competing interest.

## References

Abdullah, M.I. and Fredriksen, S. (2004) Production, respiration and exudation of dissolved organic matter by the kelp *Laminaria hyperborea* along the west coast of Norway. J Mar Biol Assoc UK 84: 887–894.

Abreu, A.C.I., Bello, M.D., Bunse, C., and Pinhassi, J. (2022) Warmer temperatures favor slower-growing bacteria in natural marine communities. bioRxiv. doi: 10.1101/2022.07.13.499956

Alonso-Sáez, L., Díaz-Pérez, L., and Morán, X.A.G. (2015) The hidden seasonality of the rare biosphere in coastal marine bacterioplankton. Environ Microbiol 17: 3766–3780.

Alonso, C., Warnecke, F., Amann, R., and Pernthaler, J. (2007) High local and global diversity of Flavobacteria in marine plankton. Environ Microbiol 9: 1253–1266.

Van Alstyne, K.L., McCarthy, J.J., Hustead, C.L., and Kearns, L.J. (1999) Phlorotannin allocation among tissues of northeastern Pacific kelps and rockweeds. J Phycol 35: 483– 492.

Ar Gall, E., Kupper, F.C., and Kloareg, B. (2004) A survey of iodine content in *Laminaria digitata*. Bot Mar 47: 30–37.

Barbeyron, T., L’Haridon, S., Corre, E., Kloareg, B., and Potin, P. (2001) *Zobellia galactanovorans* gen. nov., sp. nov., a marine species of Flavobacteriaceae isolated from a red alga, and classification of [Cytophaga] uliginosa (ZoBell and Upham 1944) Reichenbach 1989 as Zobellia uliginosa gen. nov., comb. nov. Int J Syst Evol Microbiol 51: 985–97.

Barbeyron, T., Thiébaud, M., Le Duff, N., Martin, M., Corre, E., Tanguy, G., et al. (2021) *Zobellia roscoffensis* sp. nov. and *Zobellia nedashkovskayae* sp. nov., two flavobacteria from the epiphytic microbiota of the brown alga *Ascophyllum nodosum*, and emended description of the genus *Zobellia*. Int J Syst Evol Microbiol 71: doi: 10.1099/ijsem.0.004913.

Barbeyron, Tristan, Thomas, F., Barbe, V., Teeling, H., Schenowitz, C., Dossat, C., et al. (2016) Habitat and taxon as driving forces of carbohydrate catabolism in marine heterotrophic bacteria: Example of the model algae-associated bacterium *Zobellia galactanivorans* Dsij^T^. Environ Microbiol 18: 4610–4627.

Bartlau, N., Wichels, A., Krohne, G., Adriaenssens, E.M., Heins, A., Fuchs, B.M., et al. (2022) Highly diverse flavobacterial phages isolated from North Sea spring blooms. ISME J 16: 555–568.

Bengtsson, M.M., Sjøtun, K., and Øvreås, L. (2010) Seasonal dynamics of bacterial biofilms on the kelp *Laminaria hyperborea*. Aquat Microb Ecol 60: 71–83.

Brunet, M., de Bettignies, F., Le Duff, N., Tanguy, G., Davoult, D., Leblanc, C., et al. (2021) Accumulation of detached kelp biomass in a subtidal temperate coastal ecosystem induces succession of epiphytic and sediment bacterial communities. Environ Microbiol 23: 1638–1655.

Brunet, M., Le Duff, N., Barbeyron, T., and Thomas, F. (2022) Consuming fresh macroalgae induces specific catabolic pathways, stress reactions and Type IX secretion in marine flavobacterial pioneer degraders. ISME J 16: 2027–2039.

Brunet, Maéva, Le Duff, N., Fuchs, B.M., Amann, R., Barbeyron, T., and Thomas, F. (2021) Specific detection and quantification of the marine flavobacterial genus *Zobellia* on macroalgae using novel qPCR and CARD-FISH assays. Syst Appl Microbiol 44: 126269.

Buchan, A., LeCleir, G.R., Gulvik, C.A., and González, J.M. (2014) Master recyclers: features and functions of bacteria associated with phytoplankton blooms. Nat Rev Microbiol 12: 686–698.

Bunse, C., Koch, H., Breider, S., Simon, M., and Wietz, M. (2021) Sweet spheres: Succession and CAZyme expression of marine bacterial communities colonising a mix of alginate and pectin particles . Environ Microbiol 23: 3130-3148.

Bunse, C. and Pinhassi, J. (2017) Marine Bacterioplankton Seasonal Succession Dynamics. Trends Microbiol 25: 494–505.

Campbell, B.J., Yu, L., Heidelberg, J.F., and Kirchman, D.L. (2011) Activity of abundant and rare bacteria in a coastal ocean. Proc Natl Acad Sci.

Caracciolo, M., Rigaut-Jalabert, F., Romac, S., Mahé, F., Forsans, S., Gac, J.P., et al. (2022) Seasonal dynamics of marine protist communities in tidally mixed coastal waters. Mol Ecol 31: 3761–3783.

Chernysheva, N., Bystritskaya, E., Stenkova, A., Golovkin, I., Nedashkovskaya, O., and Isaeva, M. (2019) Comparative genomics and CAZyme genome repertoires of marine *Zobellia amurskyensis* KMM 3526^T^ and *Zobellia laminariae* KMM 3676^T^. Mar Drugs 17: 661.

Cocquempot, L., Delacourt, C., Paillet, J., Riou, P., Aucan, J., Castelle, B., et al. (2019) Coastal ocean and nearshore observation: A French case study. Front Mar Sci 6: 1–17.

Díez-Vives, C., Nielsen, S., Sánchez, P., Palenzuela, O., Ferrera, I., Sebastián, M., et al. (2019) Delineation of ecologically distinct units of marine *Bacteroidetes* in the Northwestern Mediterranean Sea. Mol Ecol 28: 2846–2859.

Dogs, M., Wemheuer, B., Wolter, L., Bergen, N., Daniel, R., Simon, M., and Brinkhoff, T. (2017) *Rhodobacteraceae* on the marine brown alga *Fucus spiralis* are abundant and show physiological adaptation to an epiphytic lifestyle. Syst Appl Microbiol 40: 370– 382.

Drebes, G., Kühn, S.F., Gmelch, A., and Schnepf, E. (1996) *Cryothecomonas aestivalis* sp. nov., a colourless nanoflagellate feeding on the marine centric diatom Guinardia delicatula (Cleve) Hasle. Helgolander Meeresuntersuchungen 50: 497–515.

Egan, S., Harder, T., Burke, C., Steinberg, P., Kjelleberg, S., and Thomas, T. (2013) The seaweed holobiont: understanding seaweed–bacteria interactions. FEMS Microbiol Rev 37: 462–476.

Enke, T.N., Datta, M.S., Schwartzman, J., Cermak, N., Schmitz, D., Barrere, J., et al. (2019) Modular Assembly of Polysaccharide-Degrading Marine Microbial Communities. Curr Biol 29: 1528–1535.

Ficko-Blean, E., Préchoux, A., Thomas, F., Rochat, T., Larocque, R., Zhu, Y., et al. (2017) Carrageenan catabolism is encoded by a complex regulon in marine heterotrophic bacteria. Nat Commun 8:.

Florez, J.Z., Camus, C., Hengst, M.B., Marchant, F., and Buschmann, A.H. (2019) Structure of the epiphytic bacterial communities of *Macrocystis pyrifera* in localities with contrasting nitrogen concentrations and temperature. Algal Res 44: 101706.

Fournier, J.B., Rebuffet, E., Delage, L., Grijol, R., Meslet-Cladière, L., Rzonca, J., et al. (2014) The vanadium iodoperoxidase from the marine *Flavobacteriaceae* species *Zobellia galactanivorans* reveals novel molecular and evolutionary features of halide specificity in the vanadium haloperoxidase enzyme family. Appl Environ Microbiol 80: 7561–7573.

Gac, J.P., Marrec, P., Cariou, T., Guillerm, C., Macé, É., Vernet, M., and Bozec, Y. (2020) Cardinal Buoys: An Opportunity for the Study of Air-Sea CO2 Fluxes in Coastal Ecosystems. Front Mar Sci 7: 1–21.

Galand, P.E., Pereira, O., Hochart, C., Auguet, J.C., and Debroas, D. (2018) A strong link between marine microbial community composition and function challenges the idea of functional redundancy. ISME J 12: 2470–2478.

Gattuso, J.-P., Frankignoulle, M., and Wollast, R. (1998) Carbon and carbonate metabolism in coastal aquatic ecosystems. Annu Rev Ecol Syst 29: 405–434.

Gilbert, J.A., Field, D., Swift, P., Newbold, L., Oliver, A., Smyth, T., et al. (2009) The seasonal structure of microbial communities in the Western English Channel. Environ Microbiol 11: 3132–3139.

Del Giorgio, P.A. and Cole, J.J. (1998) Bacterial growth efficiency in natural aquatic systems. Annu Rev Ecol Syst 29: 503–541.

Goberville, E., Beaugrand, G., Sautour, B., and Tréguer, P. (2010) Climate-driven changes in coastal marine systems of western Europe. Mar Ecol Prog Ser 408: 129–147.

Gobet, A., Barbeyron, T., Matard-Mann, M., Magdelenat, G., Vallenet, D., Duchaud, E., and Michel, G. (2018) Evolutionary evidence of algal polysaccharide degradation acquisition by *Pseudoalteromonas carrageenovora* 9^T^ to adapt to macroalgal niches. Front Microbiol 9: 1–16.

Goecke, F.R., Labes, A., Wiese, J., and Imhoff, J.F. (2010) Chemical interactions between marine macroalgae and bacteria. Mar Ecol Prog Ser.

Grigorian, Eugénie, Groisillier, A., Thomas, F., Leblanc, C., and Walsh, D.A. (2021) Functional Characterization of a L-2-Haloacid Dehalogenase From *Zobellia galactanivorans* Dsij^T^ Suggests a Role in Haloacetic Acid Catabolism and a Wide Distribution in Marine Environments. Front Microbiol 12: 1–15.

Grigorian, E., Roret, T., Czjzek, M., Leblanc, C., and Delage, L. (2023) X-ray structure and mechanism of ZgHAD , a L -2-haloacid dehalogenase from the marine flavobacterium *Zobellia galactanivorans*. Protein Sci 32: 1–12.

Groisillier, A., Labourel, A., Michel, G., and Tonon, T. (2015) The mannitol utilization system of the marine bacterium *Zobellia galactanivorans*. Appl Environ Microbiol 81: 1799–1812.

Guillou, L., Viprey, M., Chambouvet, A., Welsh, R.M., Kirkham, A.R., Massana, R., et al. (2008) Widespread occurrence and genetic diversity of marine parasitoids belonging to Syndiniales (Alveolata). Environ Microbiol 10: 3349–3365.

Hahnke, R.L. and Harder, J. (2013) Phylogenetic diversity of Flavobacteria isolated from the North Sea on solid media. Syst Appl Microbiol 36: 497–504.

Harms, H., Klöckner, A., Schrör, J., Josten, M., Kehraus, S., Crüsemann, M., et al. (2018) Antimicrobial Dialkylresorcins from Marine-Derived Microorganisms: Insights into Their Mode of Action and Putative Ecological Relevance. Planta Med.

Heins, A. and Harder, J. (2023) Particle-associated bacteria in seawater dominate the colony- forming microbiome on ZoBell marine agar. FEMS Microbiol Ecol 99: 1–11.

Huang, G., Vidal-Melgosa, S., Sichert, A., Becker, S., Fang, Y., Niggemann, J., et al. (2021) Secretion of sulfated fucans by diatoms may contribute to marine aggregate formation. Limnol Oceanogr 66: 3768–3782.

Ihua, M.W., FitzGerald, J.A., Guiheneuf, F., Jackson, S.A., Claesson, M.J., Stengel, D.B., and Dobson, A.D.W. (2020) Diversity of bacteria populations associated with different thallus regions of the brown alga *Laminaria digitata*. PLoS One 15: 1–15.

Kabisch, A., Otto, A., König, S., Becher, D., Albrecht, D., Schüler, M., et al. (2014) Functional characterization of polysaccharide utilization loci in the marine *Bacteroidetes* “*Gramella forsetii*” KT0803. ISME J 8: 1492–502.

Kirchman, D.L. (2002) The ecology of Cytophaga-Flavobacteria in aquatic environments. FEMS Microbiol Ecol 39: 91–100.

Klindworth, A., Pruesse, E., Schweer, T., Peplies, J., Quast, C., Horn, M., and Glöckner, F.O. (2013) Evaluation of general 16S ribosomal RNA gene PCR primers for classical and next-generation sequencing-based diversity studies. Nucleic Acids Res 41: 1–11.

Kloareg, B., Badis, Y., Cock, J.M., and Michel, G. (2021) Role and Evolution of the Extracellular Matrix in the Acquisition of Complex Multicellularity in Eukaryotes[: A Macroalgal Perspective. Genes (Basel) 12: 1059.

Koch, H., Dürwald, A., Schweder, T., Noriega-Ortega, B., Vidal-Melgosa, S., Hehemann, J.H., et al. (2019) Biphasic cellular adaptations and ecological implications of *Alteromonas macleodii* degrading a mixture of algal polysaccharides. ISME J 13: 92– 103.

Küpper, F.C., Kloareg, B., Guern, J., and Potin, P. (2001) Oligoguluronates elicit an oxidative burst in the brown algal kelp *Laminaria digitata*. Plant Physiol 125: 278–291.

Lami, R., Grimaud, R., Sanchez-Brosseau, S., Six, C., Thomas, F., West, N.J., et al. (2021) Marine bacterial models for experimental biology. In Established and emerging marine organisms in experimental biology.

Lankiewicz, T.S., Cottrell, M.T., and Kirchman, D.L. (2016) Growth rates and rRNA content of four marine bacteria in pure cultures and in the Delaware estuary. ISME J 10: 823– 832.

Lauro, F.M., McDougald, D., Thomas, T., Williams, T.J., Egan, S., Rice, S., et al. (2009) The genomic basis of trophic strategy in marine bacteria. Proc Natl Acad Sci U S A 106: 15527–15533.

Lefran, A., Hernández-Fariñas, T., Gohin, F., and Claquin, P. (2021) Decadal trajectories of phytoplankton communities in contrasted estuarine systems in an epicontinental sea. Estuar Coast Shelf Sci 258:.

Martin, M., Portetelle, D., and Michel, G. (2014) Microorganisms living on macroalgae[: diversity , interactions , and biotechnological applications. Appl Microbiol Biotechnol 98: 2917–2935.

Massana, R., Unrein, F., Rodríguez-Martínez, R., Forn, I., Lefort, T., Pinhassi, J., and Not, F. (2009) Grazing rates and functional diversity of uncultured heterotrophic flagellates. ISME J 3: 588–595.

Mensch, B., Neulinger, S.C., Künzel, S., Wahl, M., and Schmitz, R.A. (2020) Warming, but Not Acidification, Restructures Epibacterial Communities of the Baltic Macroalga *Fucus vesiculosus* With Seasonal Variability. Front Microbiol 11: 1–18.

Miranda, L.N., Hutchison, K., Grossman, A.R., and Brawley, S.H. (2013) Diversity and abundance of the bacterial community of the red macroalga *Porphyra umbilicalis*: Did bacterial farmers produce macroalgae? PLoS One 8: e58269.

Myklestad, S.M. (1995) Release of extracellular products by phytoplankton with special emphasis on polysaccharides. Sci Total Environ 165: 155–164.

Nedashkovskaya, O., Otstavnykh, N., Zhukova, N., Guzev, K., Chausova, V., Tekutyeva, L., et al. (2021) *Zobellia barbeyronii* sp. nov., a New Member of the Family *Flavobacteriaceae*, Isolated from Seaweed, and Emended Description of the Species *Z. amurskyensis*, *Z. laminariae*, Z. russellii and Z. uliginosa. Diversity 13: 520.

Nedashkovskaya, O., Suzuki, M., Vancanneyt, M., Cleenwerck, I., Lysenko, A., Mikhailov, V., and Swings, J. (2004) *Zobellia amurskyensis* sp nov., Zobellia laminariae sp nov and Zobellia russellii sp nov., novel marine bacteria of the family Flavobacteriaceae. International J Syst Evol Microbiol 54: 1643–1648.

Nedashkovskaya, O.I. and Suzuki, M. (2015) Zobellia. In Bergey’s Manual of Systematics of Archaea and Bacteria. M.E. Trujillo, S. Dedysh, P. DeVos, B. Hedlund, P. Kämpfer, F.A.R. and W.B.W. (ed). John Wiley & Sons, Inc.

Nitschke, U., Walsh, P., McDaid, J., and Stengel, D.B. (2018) Variability in iodine in temperate seaweeds and iodine accumulation kinetics of *Fucus vesiculosus* and *Laminaria digitata* (Phaeophyceae, Ochrophyta). J Phycol 54: 114–125.

Paix, B., Layglon, N., Poupon, C. Le, Onofrio, S.D., Misson, B., and Garnier, C. (2021) Integration of spatio-temporal variations of surface metabolomes and epibacterial communities highlights the importance of copper stress as a major factor shaping host- microbiota interactions within a Mediterranean seaweed holobiont. Microbiome 9: 201.

Paix, B., Othmani, A., Debroas, D., Culioli, G., and Briand, J.F. (2019) Temporal covariation of epibacterial community and surface metabolome in the Mediterranean seaweed holobiont *Taonia atomaria*. Environ Microbiol. 21: 3346–3363.

Pedler, B.E., Aluwihare, L.I., and Azam, F. (2014) Single bacterial strain capable of significant contribution to carbon cycling in the surface ocean. Proc Natl Acad Sci U S A 111: 7202–7207.

R Core Team (2018) R: A Language and Environment for Statistical Computing.

Ramirez-Puebla, S.T., Weigel, B.L., Jack, L., Schlundt, C., Pfister, C.A., and Welch, Mark, J.L. (2022) Spatial organization of the kelp microbiome at micron scales. Microbiome 10: 52.

Reisky, L., Préchoux, A., Zühlke, M.-K., Bäumgen, M., Robb, C.S., Gerlach, N., et al. (2019) A complex enzyme cascade degrades the polysaccharide ulvan from green algae. Nat Chem Biol 15: 803–812.

Röttjers, L. and Faust, K. (2018) From hairballs to hypotheses–biological insights from microbial networks. FEMS Microbiol Rev 42: 761–780.

Salaün, S., La Barre, S., Dos Santos-Goncalvez, M., Potin, P., Haras, D., and Bazire, A. (2012) Influence of exudates of the kelp *Laminaria digitata* on biofilm formation of associated and exogenous bacterial epiphytes. Microb Ecol 64: 359–69.

Santos, I.R., Hatje, V., Serrano, O., Bastviken, D., and Krause-Jensen, D. (2022) Carbon sequestration in aquatic ecosystems: Recent advances and challenges. Limnol Oceanogr 67: S1–S5.

Sichert, A., Corzett, C.H., Schechter, M.S., Unfried, F., Markert, S., Becher, D., et al. (2020) Verrucomicrobia use hundreds of enzymes to digest the algal polysaccharide fucoidan. Nat Microbiol.

Singh, R.P. and Reddy, C.R.K. (2016) Unraveling the functions of the macroalgal microbiome. Front Microbiol 6: 1–8.

Staufenberger, T., Thiel, V., Wiese, J., and Imhoff, J.F. (2008) Phylogenetic analysis of bacteria associated with *Laminaria saccharina*. FEMS Microbiol Ecol 64: 65–77.

Takemura, A.F., Chien, D.M., and Polz, M.F. (2014) Associations and dynamics of *Vibrionaceae* in the environment, from the genus to the population level. Front Microbiol 5: 1–26.

Teeling, H., Fuchs, B.M., Becher, D., Klockow, C., Gardebrecht, A., Bennke, C.M., et al. (2012) Substrate-controlled succession of marine bacterioplankton populations induced by a phytoplankton bloom. Science 336: 608–11.

Teeling, H., Fuchs, B.M., Bennke, C.M., Krüger, K., Chafee, M., Kappelmann, L., et al. (2016) Recurring patterns in bacterioplankton dynamics during coastal spring algae blooms. Elife 5: 1–29.

Thomas, F., Barbeyron, T., and Michel, G. (2011) Evaluation of reference genes for real-time quantitative PCR in the marine flavobacterium *Zobellia galactanivorans*. J Microbiol Methods 84: 61–66.

Thomas, F., Barbeyron, T., Tonon, T., Génicot, S., Czjzek, M., and Michel, G. (2012) Characterization of the first alginolytic operons in a marine bacterium: from their emergence in marine *Flavobacteriia* to their independent transfers to marine *Proteobacteria* and human gut *Bacteroides*. Environ Microbiol 14: 2379–94.

Thomas, F., Bordron, P., Eveillard, D., and Michel, G. (2017) Gene Expression Analysis of *Zobellia galactanivorans* during the Degradation of Algal Polysaccharides Reveals both Substrate-Specific and Shared Transcriptome-Wide Responses. Front Microbiol 8: 1808.

Ulrich, J.F., Gräfe, M.S., Dhiman, S., Wienecke, P., Arndt, H., and Wichard, T. (2022) Thallusin Quantification in Marine Bacteria and Algae Cultures. Mar. Drugs 20: 690.

Weigel, B.L. and Pfister, C.A. (2019) Successional dynamics and seascape-level patterns of microbial communities on the canopy-forming kelps *Nereocystis luetkeana* and *Macrocystis pyrifera*. Front Microbiol 10: 1–17.

Xia, L.C., Steele, J. a, Cram, J. a, Cardon, Z.G., Simmons, S.L., Vallino, J.J., et al. (2011) Extended local similarity analysis (eLSA) of microbial community and other time series data with replicates. BMC Syst Biol 5: S15.

